# A high-resolution blood immune cell doublet atlas via imaging spectral cytometry

**DOI:** 10.64898/2026.09.03.749150

**Authors:** Cheryl Kim, Denise Hinz, Aaron Middlebrook, Aaron J Tyznik, Louise M D’Cruz, Thomas Scriba, Bjoern Peters, Julie G Burel

## Abstract

Circulating cell-cell complexes (doublets) provide critical insights into in vivo immune interactions. Here, we leverage high-dimensional imaging spectral cytometry to systematically map the landscape of immune doublets in human peripheral blood. By integrating morphometric imaging features with a 22-color spectral panel, we established a gating workflow that cleanly segregates genuine, physically interacting complexes from coincidental transits and platelet contaminants, and resolved 12 highly pure homotypic and heterotypic immune doublet configurations in peripheral blood mononuclear cells (PBMC). At steady state, each doublet population displayed unique abundance, affinity and surface phenotypic profiles relative to circulating singlets. Principal component analysis (PCA) of quantitative imaging features further revealed lineage-specific topologies distinguishing singlet cells, homotypic doublets, and heterotypic pairs. Finally, comparative cohort analysis demonstrated that while doublet frequencies, affinities and surface phenotypes remain comparable between healthy donors and patients with tuberculosis (TB) disease, imaging-derived features uncovered disease-specific alterations in doublets morphology. Specifically, high-dimensional clustering using imaging parameters showed that TB disease significantly enriches for a tightly synapsed T cell-monocyte cluster characterized by high CD3-CD14 pixel correlation. Collectively, this study establishes a robust framework for mapping circulating cell-cell networks and demonstrates the superior sensitivity of imaging features over conventional fluorescence for capturing functional immune interactions in health and disease.

## Introduction

While cell doublets are routinely observed in flow cytometry, they are historically discarded as technical artifacts resulting from *ex vivo* sample manipulation. However, emerging evidence demonstrates that a significant fraction of immune cell doublets detected in human peripheral blood represent genuine, biologically functional interactions (1). For instance, we previously showed that circulating T cell-monocyte doublets exhibit polarization of adhesion molecules at the point of contact (2,3) and possess transcriptomic profiles distinct from singlets during infection, characterized by the upregulation of MHC class II, cytotoxicity, and interferon signaling pathways (3). Similarly, independent groups have identified unique T cell-monocyte doublet transcriptomic signatures in individuals with COVID-19 (4) and HIV (5), which correlate with elevated MHC-II expression, heightened immune activation, and – in the case of HIV – higher viral loads.

To date, studies of circulating doublets have focused primarily on pairings between T cells and monocytes, the two most abundant immune cell subsets in blood. However, flow cytometry data of human peripheral blood mononuclear cells (PBMC) reveal the presence of events co-expressing a combination of T cell, B cell, NK cell, and myeloid cell surface markers, suggesting a far wider diversity of immune complexes in circulation. For example, we previously demonstrated that the majority of CD3^+^CD19^+^ live “singlet” events in human PBMC are physical T cell-B cell complexes rather than dual-expressing single cells (6). Non-T cell doublets have also been identified via single-cell transcriptomics, including myeloid-B cell complexes in the lung, ileum and spleen of healthy individuals (7), and myeloid-NK cell complexes in blood during COVID-19 convalescence (8). These various cell-cell combinations are expected to serve important immune functions (9–11).

Real-time imaging spectral cytometry represents a major technological leap, combining high-dimensional spectral fluorescence with real-time morphometric image acquisition and reconstruction (12,13). This innovation allows for much larger panels than those used in traditional imaging cytometry. Crucially, real-time image processing enables high-throughput sorting based on active imaging parameters – a significant advancement over traditional, post-acquisition pixel-based imaging cytometry. Key immunological applications for this technology include high-resolution cell cycle analysis, nuclear translocation, and cell-cell interactions.

Here, we present the first comprehensive atlas of circulating immune cell doublets in human blood using real-time imaging spectral cytometry. Utilizing a 22-color spectral panel spanning all major PBMC lineages, we designed a multi-level gating strategy that integrates high-dimensional fluorescence with morphometric imaging features to systematically resolve 12 highly pure homotypic and heterotypic doublet configurations. We mapped their distinct abundance, affinity, and phenotypic profiles at steady state, and demonstrate that under pathological conditions, such as active tuberculosis (TB), morphometric imaging parameters reveal disease-specific signatures that are missed by traditional fluorescence intensity profiling alone. This study establishes a robust, validated methodological resource to guide future high-fidelity investigations of circulating immune complexes in health and disease.

## Results

### Development and optimization of a 22-color spectral imaging panel for circulating doublet profiling

We designed a 22-color spectral immunophenotyping panel targeting key lineage markers for all major immune cell lineages present in human PBMC (**Table 1**). The three dedicated fluorescent imaging detectors were assigned to identify the most abundant circulating lineages – T cells, B cells, and CD14^+^ monocytes – using antibodies against CD3, CD19 and CD14, respectively. The remaining 19 non-imaging fluorescent channels incorporated a viability dye and antibodies targeting NK cells, dendritic cells (DCs), platelets, and leukocytes, alongside differentiation and activation markers to comprehensively profile sub-lineages. Overall panel complexity was computationally evaluated using the Complexity Score (or Index) in BD^®^ Research Cloud, yielding a manageable score of 7.35 (**Figure 1A**).

**Figure 1:**
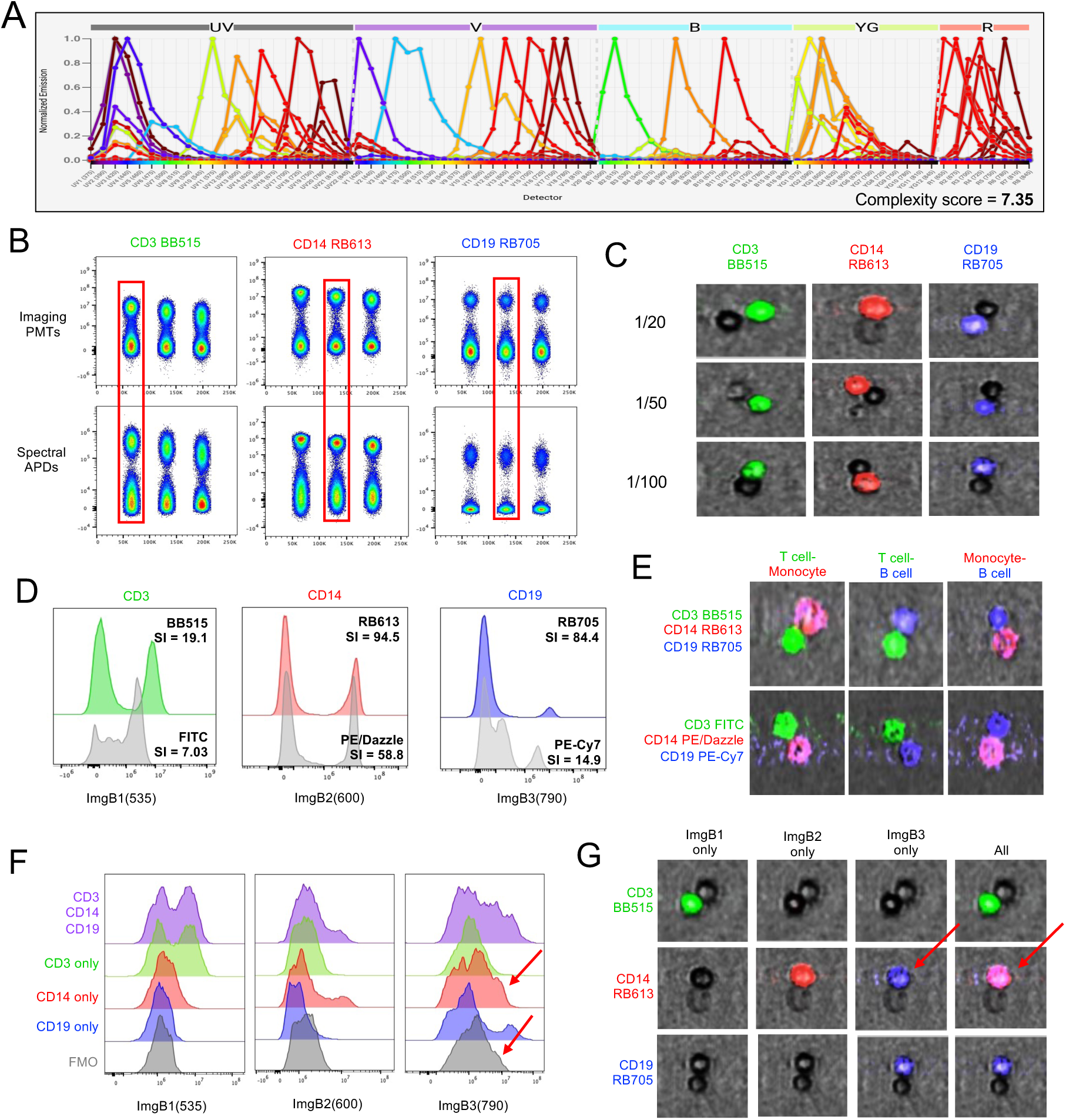
Optimization of a 22-color spectral imaging panel for circulating doublet profiling on the BD FACS Discover^TM^ S8. (A) Full spectral signature and complexity score of the 22-color panel evaluated using the BD Research Cloud. (B) Co-titration of CD3 BB515, CD14 RB613 and CD19 RB705 antibodies (1:20, 1:50 and 1:100 dilution) in human PBMC. Respective stain indexes for singlets were calculated in both imaging detectors (PMTs, top row) and spectral detectors (APDs, bottom row). (C) Representative BD FACSChorus™ image wall galleries of the co-titration in CD3^+^, CD14^+^ or CD19^+^ synaptic doublets (as gated in Figure 2C) overlaying the brightfield (LightLoss) and the corresponding single channel. Stain indexes of newer generation dyes (CD3 BB515, CD14 RB613, CD19 RB705) compared to legacy dyes (CD3 FITC, CD14 PE/Dazzle, CD19 PE-Cy7) across (D) imaging PMTs and (E) representative images from the image wall. Imaging channel overlap assessment using single-color, fluorescence-minus-one (FMO), and fully stained controls via (F) histogram overlays and (G) image wall galleries. Synaptic doublets were gated as shown in Figure 2C and displayed in the BD FACSChorus image wall selecting each individual imaging channel and all 3 imaging channels combined (All; CD3, green; CD14, red; CD19, blue). Red arrows indicate the spillover of CD14 into ImgB3. Data derived from healthy PBMC (n=1).

**Table 1:**
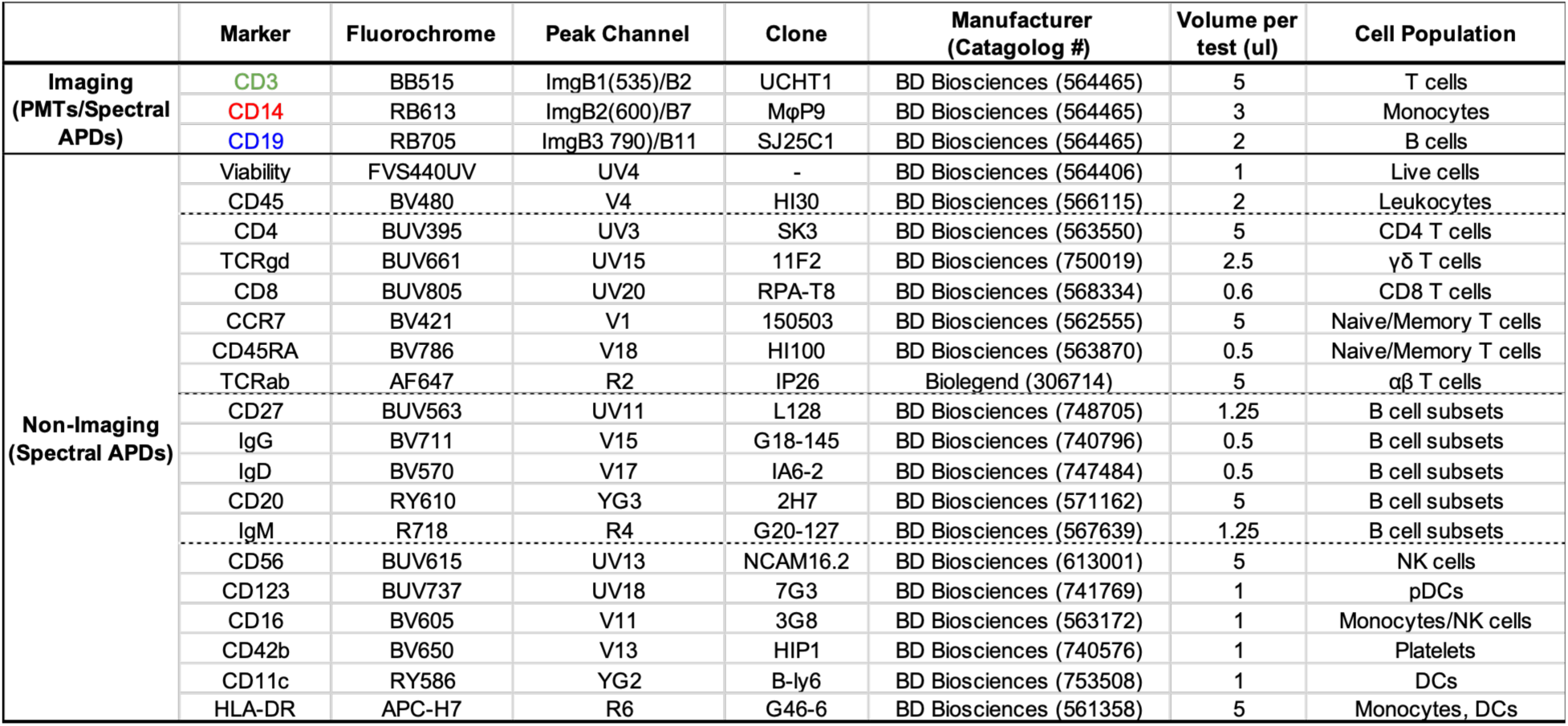
A 22-color spectral and imaging panel for circulating doublet profiling.

The panel was designed and optimized for the FACSDiscover^TM^ S8 spectral imaging cell sorter operating on BD FACSChorus^TM^ version 5.4. In this software version, fluorescence compensation and spectral unmixing were not available for imaging detectors. To maximize signal-to-noise ratios and minimize spectral overlap across the three imaging detectors, we conducted a coordinated titration of CD3, CD14, and CD19 imaging antibodies (14). Optimal concentrations were established based on stain index, brightness and reduced background in the Image Wall: CD3 Brilliant Blue™ 515 (BB515) was selected at 1:20 dilution, CD14 Real Blue™ 613 (RB613) at 1:50, and CD19 Real Blue™ 705 (RB705) at 1:50 (**Figures 1B**, **1C**). Direct benchmarking against legacy fluorophores (CD3 FITC, CD14 PE/Dazzle, and CD19 PE-Cy7) confirmed that next-generation RB and BB dyes offered superior brightness with significantly reduced background noise and spillover spreading driven by minimal cross-laser excitation (**Figures 1D**, **1E**). Finally, fluorescence-minus-one (FMO) controls were evaluated to quantify detector spillover and set gating boundaries. While CD3 BB515 and CD19 RB705 displayed negligible spillover into adjacent imaging channels, CD14 RB613 signal was detected within the ImgB3 channel across both CD14 single-color and CD19 FMO controls conditions (**Figures 1F, 1G**). Consequently, CD14 expression manifests as a composite signal from both ImgB2 and ImgB3 detectors in the ImageWall display settings (magenta color, **Figure 1G**). While not a problem for fluorescence and single imaging parameters, this optical artifact artificially inflates spatial overlap metrics between CD14 and CD19, thus dual-lineage CD14/CD19 spatial parameters (such as pixel correlation and delta center of mass) were excluded from downstream analyses.

### Marker-independent morphometric gating isolates physically interacting doublets

Next, we established a gating strategy to identify cell-cell doublet populations using morphometric parameters derived exclusively from imaging detectors – specifically Side Scatter (SSC) imaging and LightLoss imaging (analogous to brightfield in traditional imaging cytometry). Because this gating scheme relies solely on global imaging parameters rather than individual markers, it is universally applicable to any panel acquired on the FACSDiscover S8 instrument. Starting from the viable CD45^+^ leukocyte population, we combined radial moment and eccentricity metrics from the SSC imaging detector to segregate doublets from singlets (**Figure 2A**). Visual inspection of the resulting image galleries revealed heterogeneity within this preliminary gate, including residual singlets, coincidental transits, and cellular triplets (**Figure 2B**, Population i). To eliminate these contaminants, we refined the selection using LightLoss imaging-derived parameters – specifically eccentricity and radial moment (**Figures 2A**, **2B,** Population ii). Finally, we filtered for intermediate-sized doublet events by cross-referencing SSC imaging (Size) against LightLoss imaging (Area) (**Figure 2A**, Population iii). This multi-step, marker-independent morphometric pipeline yielded a highly refined population enriched for genuine, physically interacting cell-cell doublets (**Figure 2B**, Population iii).

**Figure 2:**
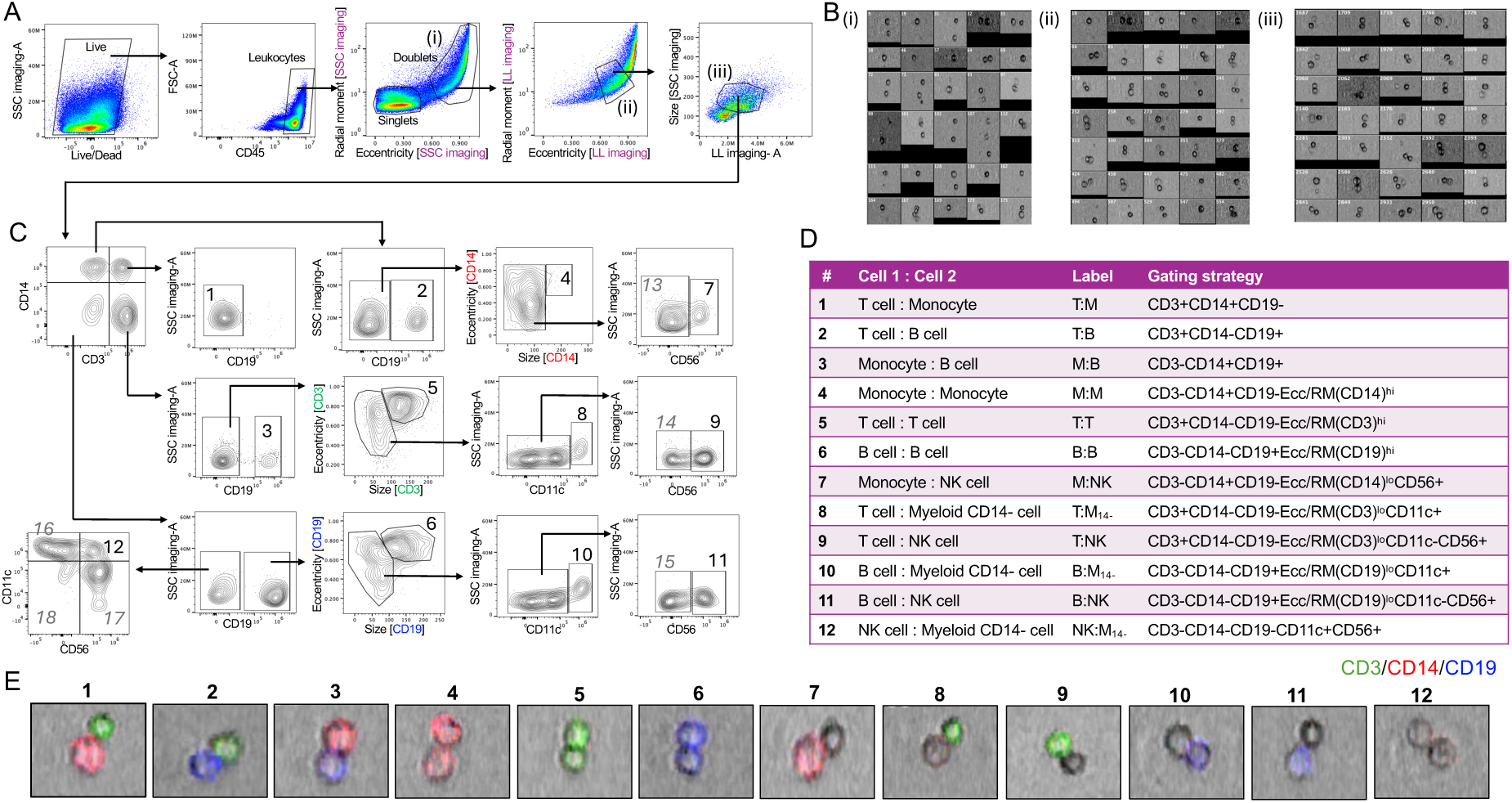
Fluorescence- and image-guided gating resolves 12 distinct physically interacting immune doublet populations. (A) Sequential, marker-independent morphometric gating strategy to identify physically interacting doublets from human PBMC (population iii) based on SSC and LightLoss imaging parameters. (B) Representative image wall galleries illustrating progressive interacting doublet enrichment from Populations i to iii. (C) Lineage gating strategy to identify homotypic and heterotypic immune doublet populations from the physically interacting doublet population iii based on fluorescence and imaging parameters. (D) Composition, label, gating definition and (E) representative composite images of the 12 validated immune doublet populations (Populations 1 to 12; CD3, green; CD14, red; CD19, blue). Image data were processed using FlowJo^TM^ (v10.10) and the BD CellView^TM^ Lens plugin (v2.1). Data derived from healthy PBMC (n=5).

### Fluorescence- and image-guided segmentation resolves 12 distinct homotypic and heterotypic immune doublet populations

Within the imaging-defined interacting doublet pool (**Figure 2A**, population iii), we leveraged the three fluorescence imaging detectors to identify T cell-B cell (T:B), T cell-CD14^+^ monocyte (T:M), and CD14^+^ monocyte-B cell (M:B) complexes (**Figure 2C**, Populations 1-3). Because these lineage markers were captured on imaging detectors, the spatial arrangement and dual-lineage identity of each population was easily visually verified via their matched image galleries (**Figure S1A**).

Next, from the single positive CD3, CD14, and CD19 doublet gates, we differentiated homotypic doublets from heterotypic complexes based on the radial moment and eccentricity measured within each fluorescence imaging channel. High eccentricity and elevated radial moment values within a single lineage channel – reflecting an elongated, doublet-like morphology equivalent to the brightfield SSC/LightLoss parameters – demarcated homotypic cell pairs (**Figure 2C**, Populations 4-6). Visual inspection of matched image galleries confirmed that this morphometric gating strategy selectively and accurately isolated true homotypic doublets (**Figure S1A**).

The remaining heterotypic configurations were resolved using non-imaging cell surface markers: CD11c to track CD14^-^ myeloid lineages, and CD56 to identify NK cells (**Figure 2C**, Populations 7-11). CD11c was selected over HLA-DR to identify myeloid cells to prevent cross-contamination from HLA-DR–expressing B cells and activated T cells. We also identified a distinct population co-expressing CD56 and CD11c within the CD3/CD14/CD19 triple-negative gate (**Figure 2C**, Population 12). Representative image galleries for each of these heterotypic doublet subpopulations are provided in **Figure S1A**.

The remaining events –single-positive for CD3, CD14, CD19, CD56 or CD11c without a partner lineage marker (**Figure 2C**, Populations 13–17) – represented ambiguous doublet populations. Image analysis revealed these events frequency comprised a single leukocyte in close spatial proximity to a smaller cellular entity resembling a platelet (**Figure S1A**). A similar profile was observed in the lineage-negative gate (**Figure 2C**, **Figure S1A**, Population 18). Expression of the platelet marker CD42b confirmed that Populations 13-18 harbored a remarkably high frequency of CD42b^+^ events (median range: 45.2%–85.2%) compared to the heterotypic and homotypic Populations 1-12 (median range: 5.6%–22.6%; **Figure S1B**), verifying that these ambiguous gates were heavily contaminated with platelets.

Purging these platelet-contaminated gates yielded 12 validated, highly pure immune doublet populations (**Figure 2D**; representative images in **Figure 2E**). Thus, combining a 5-color core lineage panel with morphometric parameters established a high-fidelity framework to comprehensively delineate the diversity of circulating cell-cell interactions in human PBMC.

### Steady-state profiling reveals distinct abundance and affinity profiles among doublet configurations

To evaluate the reproducibility and defining features of these distinct doublet populations, we applied our panel and gating strategy to a cohort of healthy donor PBMC samples (n=5). The 12 validated doublet populations constituted approximately two-thirds (median range: 59.7%–77.2%) of the total morphometrically gated, in-contact doublet population (**Figure 3A**), with the remaining one third comprising the ambiguous, platelet-contaminated populations 13-18 (**Figure 2C**). This finding underscores that while imaging-derived parameters provide an essential initial filter, integrating targeted cell-surface lineage markers is critical to achieve maximum doublet purity. Manual verification of image galleries across all five donors confirmed high purity for the 12 doublet populations (medians: 80%–96%, **Figure 3B**). Homotypic M:M and T:T doublets exhibited the highest purity, whereas the heterotypic M:B population demonstrated the lowest. Generally, populations with lower median purity displayed greater inter-individual variability.

**Figure 3:**
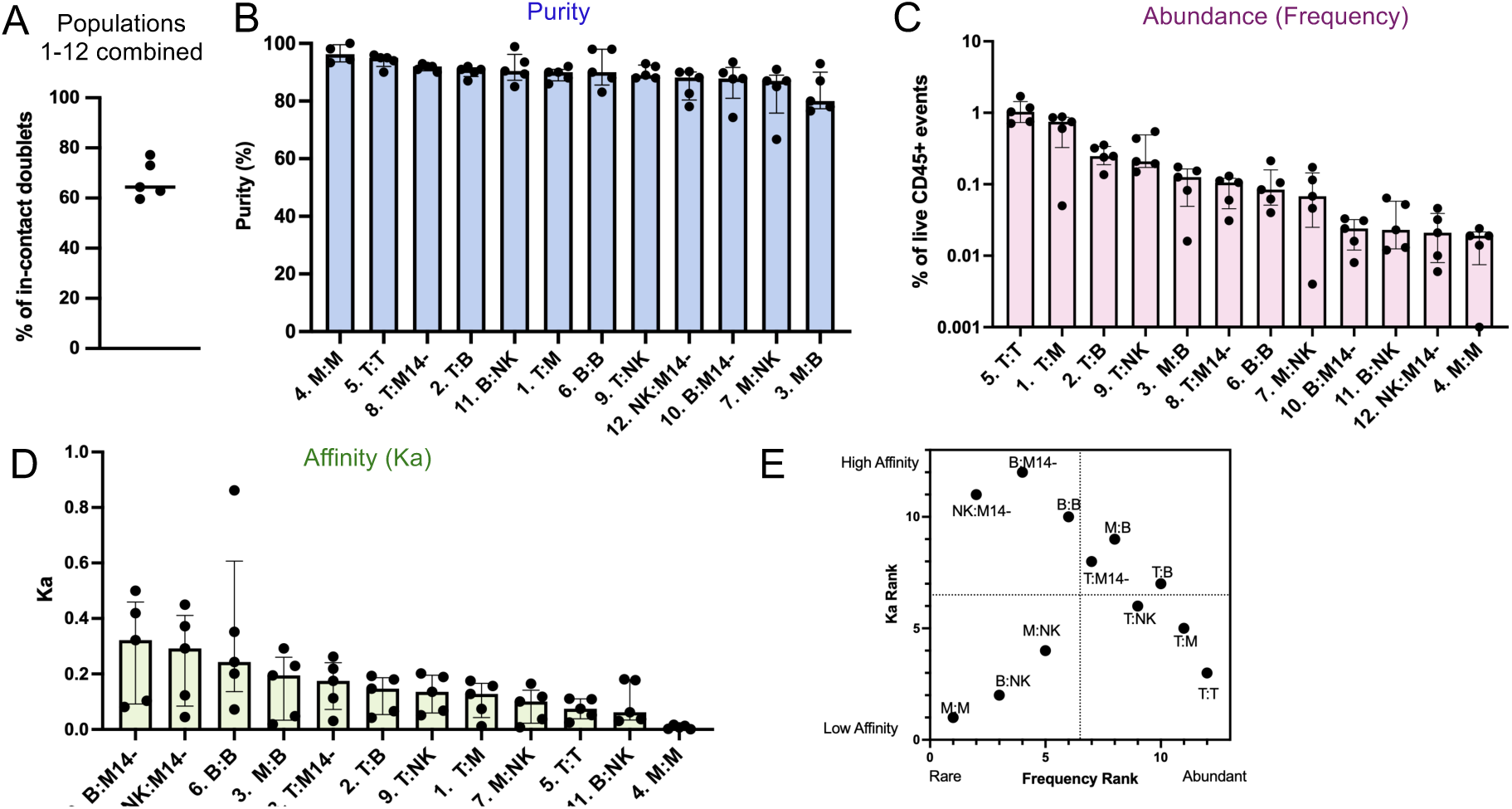
Steady-state profiling reveals distinct abundance and affinity profiles among circulating doublet configurations. (A) Cumulative frequency of the 12 validated doublet populations (Populations 1-12, Figure 2C), expressed as a percentage of physically interacting doublets (Population iii, Figure 2A). (B) Purity of the 12 validated doublet configurations determined by manual visual inspection of image galleries. (C) Frequencies (expressed as aa percentage of live CD45^+^ events) and (D) constant of association Ka of the 12 validated doublet populations, ranked by magnitude (highest to lowest). Ka was calculated by dividing doublet frequency by the product of constituent singlet frequencies, as previously reported (2). (E) Quadrant plot displaying frequency rank against Ka rank for each doublet population (Rank 1, lowest; Rank 12, highest). Individual datapoints, median and interquartile range are plotted. Image data were processed using FlowJo^TM^ (v10.10) and the BD CellView^TM^ Lens plugin (v2.1). Data derived from healthy PBMC (n=5).

Steady-state frequencies of individual doublet types varied widely, ranging from approximately 1% down to less than 0.02% of total CD45^+^ events (**Figure 3C**). T cell-containing complexes were the most prevalent, led by homotypic T:T doublets (1.03%), followed by heterotypic T:M (0.75%), T:B (0.25%), and T:NK (0.21%) pairs, whereas homotypic M:M doublets represented the rarest configuration (0.02%).

Because *ex vivo* doublet frequency is directly correlated with the baseline abundance of its constituent singlets, we calculated the constant of association Ka to mathematically normalize for precursor singlet cell frequencies (2). Indeed, singlet T cells were the most abundant population in peripheral blood, followed by monocytes (**Figures S2A**, **2B**). Heterotypic doublets containing a myeloid CD14^-^ cell exhibited the highest Ka values, displaying preferential binding toward B cells and NK cells (**Figure 3D**). Homotypic B:B doublets also demonstrated elevated Ka values, whereas homotypic T:T and M:M doublets exhibited the lowest relative affinity (**Figure 3D**).

Plotting their frequency ranking against their respective Ka ranking, all 12 doublet populations were classified into four quadrants: (i) high abundance/high affinity, (ii) high abundance/low affinity, (iii) low abundance/low affinity, and (iv) low abundance/high affinity (**Figure 3E**). Abundant, high-affinity populations included M:B, T:B, and T:M_14-_ complexes, whereas T:T, T:M and T:NK pairs fell into the abundant but low-affinity quadrant. Rare, low-affinity interactions were restricted to M:M, M:NK, and B:NK doublets, while the rare but high-affinity quadrant was populated by NK:M_14-_, B:M_14-_, and homotypic B:B pairs. Thus, decoupling cellular abundance from affinity demonstrates that circulating immune lineages do not associate stochastically, but rather exhibit distinct biological preferences for physical interaction.

### Doublet combinations display distinct surface phenotypes relative to singlets

We next leveraged the remaining 14 spectral markers from the panel to evaluate phenotypic sub-lineages within each doublet population. We first validated marker specificity by defining baseline expression across the five core singlet populations (**Figures 4A, S2C**). We confirmed three lineage-restricted T cell markers (CD4, TCRαβ, TCRγδ) and three B cell markers (CD20, IgD, IgM). Expected multi-lineage expression was also noted: CD8 was expressed on T and NK cells, while CD27 was shared across T, B, and NK lineages. CD123 was expressed on B and myeloid cells, while CD16 was shared across T, NK, and myeloid lineages. Three markers (CD45RA, CCR7, and HLA-DR) were broadly expressed by all cell types and thus excluded from this analysis. Additionally, IgG positive staining was detected on myeloid and NK cells despite pre-incubation with a commercial Fc-receptor blocking reagent, likely reflecting incomplete blocking of high-affinity IgG receptors known to be expressed in these lineages (15). A summary matrix of marker expression compatibility for each doublet configuration is presented in **Figure 4B**.

**Figure 4:**
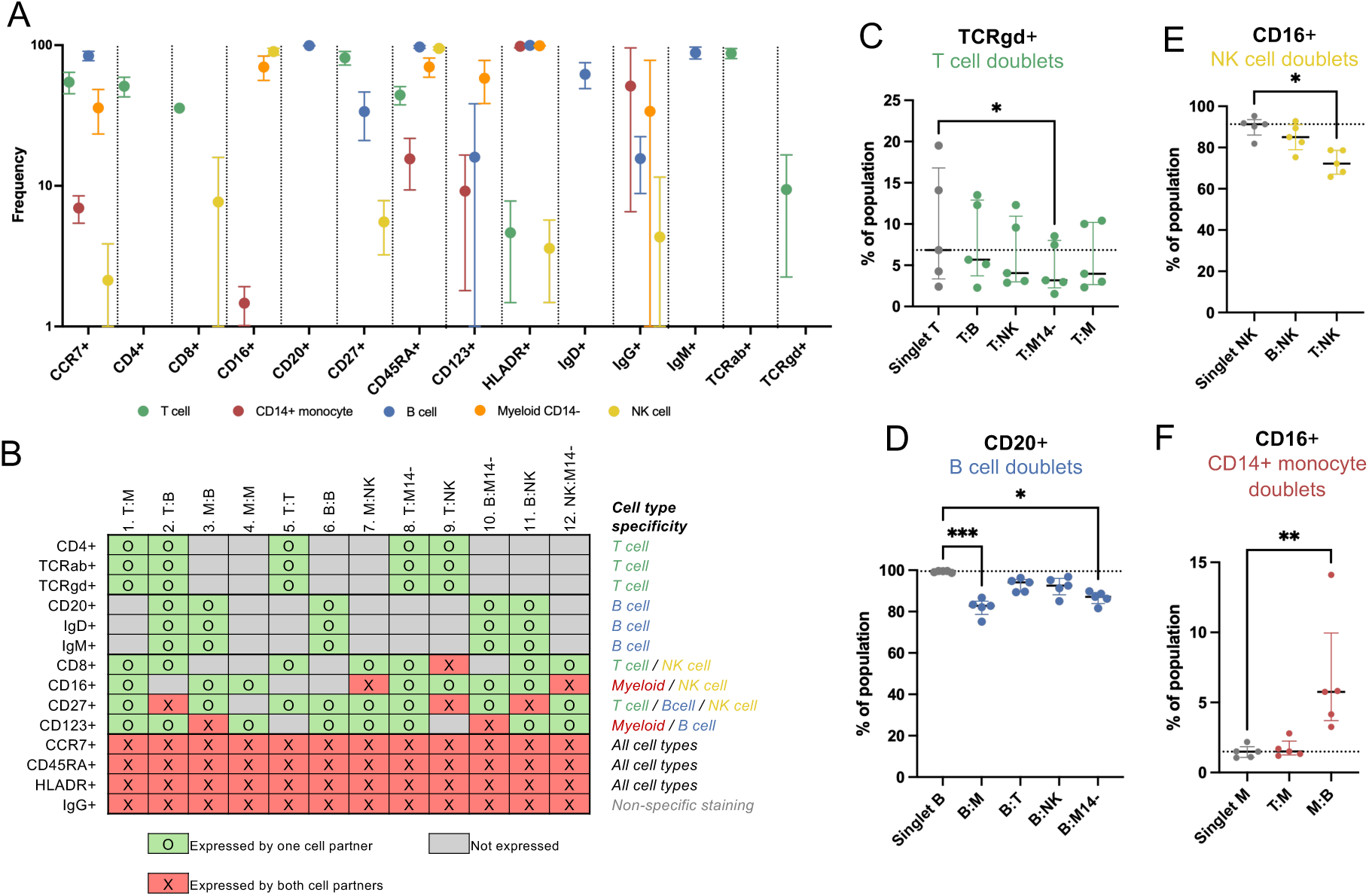
Doublet combinations display distinct surface phenotypes relative to singlets. (A) Positivity frequencies of 14 surface markers across five major singlet populations in PBMC (as gated in Figure S2A). The positive gate for each marker was set as shown for total refined singlets (Figure S2C) and median and interquartile range are plotted; frequencies lower than 1% were omitted. (B) Marker/doublet population compatibility matrix. Frequencies of (C) TCRγδ^+^ T cells, (D) CD20^+^ B cells, (E) CD16^+^ NK cells and (F) CD16^+^ CD14^+^ monocytes within indicated heterotypic doublet configurations compared to singlet counterparts. Doublet populations were identified as shown in Figure 2C and singlets as shown in Figure S2A. Individual datapoints, median and interquartile range are plotted; dashed line represent median frequency of singlets. Frequencies of marker-positive cells within heterotypic doublet populations were compared to singlets using a non-parametric unpaired Mann-Whitney test (* *p*<0.05; ** *p*<0.01; *** *p*<0.001). Data derived from healthy PBMC (n=5).

Based on this compatibility matrix, we directly compared marker positivity in individual heterotypic doublet populations against their constituent singlet counterparts (homotypic doublets were excluded from this analysis as individual cell signals cannot be deconvoluted). Within T cell-containing complexes, T:M_14-_ complexes exhibited a reduced frequency of TCRγδ^+^ T cells (**Figure 4C**), whereas CD4, CD8 and CD27 frequencies remained unchanged relative to singlets (**Figure S3A**). Among B cell complexes, B:M and B:M_14-_ doublets demonstrated a reduction in the frequency of CD20^+^ B cells (**Figure 4D**), with no alterations in CD27, CD123, IgM and IgD profiles (**Figure S3B**). For NK cell-containing complexes, T:NK doublets exhibited a decrease in CD16 positivity (**Figure 4E**), while CD8 and CD27 remained comparable to singlets (**Figure S3C**). Analysis of CD14^+^ monocyte doublets revealed that M:B were enriched for CD16^+^ intermediate monocytes (**Figure 4F**), whereas CD123 expression was equivalent across singlets and doublets (**Figure S3D**). Finally, no differences in CD16 or CD123 positivity were observed in myeloid CD14^-^ doublets (**Figure S3E**). Collectively, these data demonstrate that specific immune sub-lineages preferentially engage in circulating cell-cell interactions.

### Doublet combinations display heterogenous imaging profiles

We next investigated whether quantitative imaging-derived parameters could discriminate between distinct doublet combinations. For every doublet population containing a T cell, a CD14^+^ monocyte, or a B cell, we extracted specific features from their respective fluorescence imaging channels (CD3, CD14, and CD19): size, diffusivity, eccentricity, short-axis moment, long-axis moment, radial moment, maximum intensity, and total intensity (**Figure 5A**). Principal Component Analysis (PCA) demonstrated that across all three lineages, singlets segregated completely from all doublet populations, while homotypic doublets formed an independent cluster distinct from heterotypic configurations (**Figures 5B-5D**). This separation was driven largely by total intensity, size, maximum intensity, and diffusivity – all of which, as expected, peaked in homotypic pairs and reached minimum values in singlets (**Figures 5B, 5D**, **S4**).

**Figure 5:**
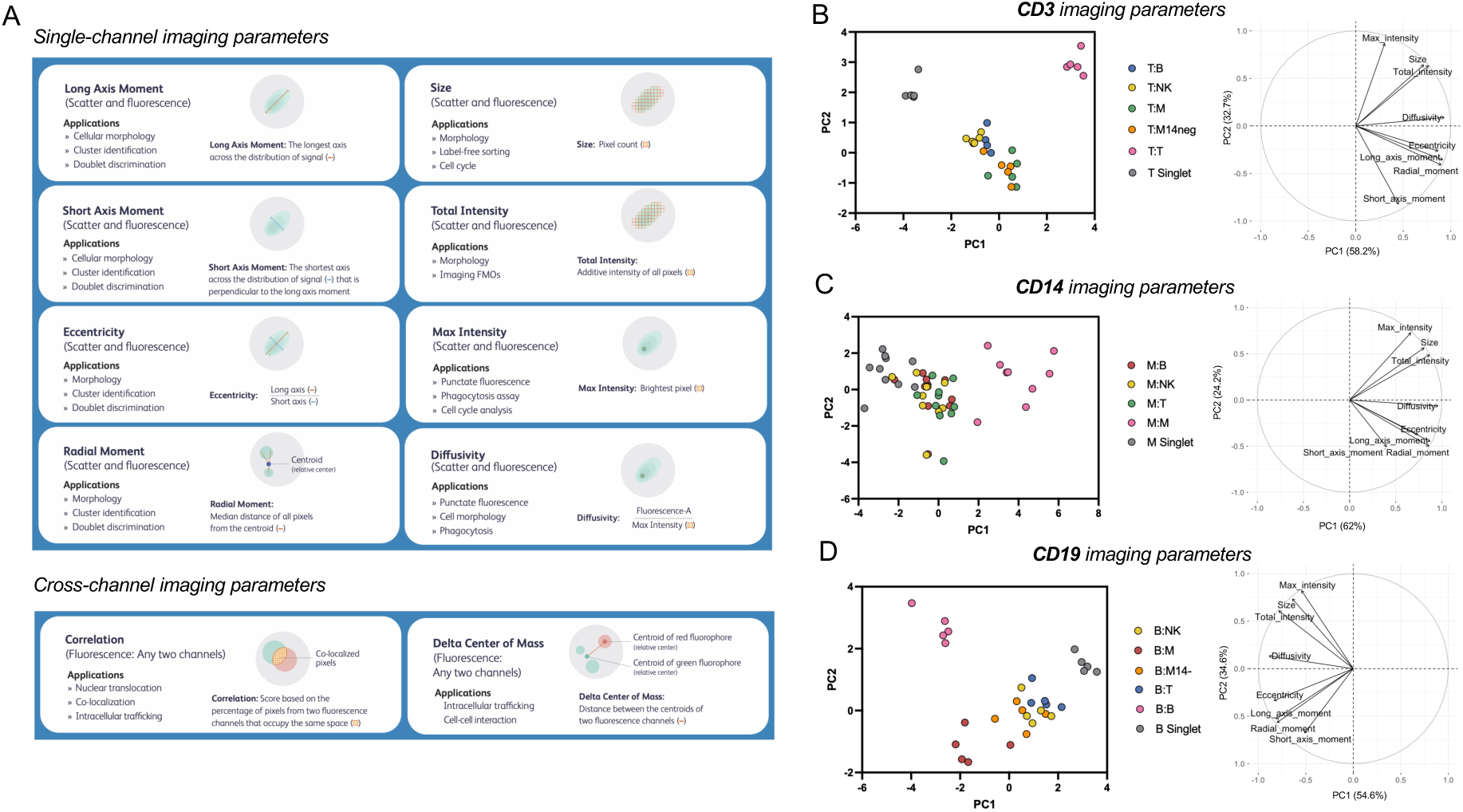
Doublet combinations display heterogenous imaging profiles. (A) Schematic representation of quantitative imaging metrics extracted from fluorescence imaging channels. PCA plots (right) and bivariate feature plots (left) of (B) CD3 imaging parameters in T cell-containing doublets, (C) CD14 imaging parameters in CD14^+^ monocyte-containing doublets and (D) CD19 imaging parameters in B cell-containing doublets. Doublet populations were identified as shown in Figure 2C and singlets as shown in Figure S2A. Data derived from healthy PBMC (n=5).

Heterotypic doublets occupied a largely overlapping morphometric space, yet displayed lineage-specific variations driven largely by short-axis moment (**Figures 5B**, **5C**). In T-cell-containing complexes, T:M and T:M_14-_ doublets separated slightly from T:B and T:NK pairs due to an increased short-axis moment (**Figure S4**). While CD14^+^ monocyte-containing heterotypic doublets remained uniform, B cell-containing complexes exhibited notable divergence: B:M doublets separated from other heterotypic B cell pairs, driven by concurrent increases in short-axis, long-axis, and radial moments (**Figure S4**). Taken together, these data demonstrate that circulating cell-cell complexes possess distinct, lineage-specific physical geometries rather than uniform morphometric profiles.

### Imaging parameters capture disease-specific doublet morphologies undetected by frequencies or surface phenotyping

Finally, we leveraged this imaging platform and gating workflow to perform a comparative analysis between healthy donors and patients with active tuberculosis (TB). We observed no statistically significant differences in the frequencies or calculated constant of association (Ka) across the 12 core doublet populations between healthy and TB cohorts (**Figures S5A**, **S5B**). Surface phenotype profiling similarly revealed minimal divergence between health and disease in T cell (**Figure S5C**), NK cell (**Figure S5E**), and myeloid CD14^-^ doublets (**Figure S5G**). Among B cell doublets, both singlets and doublets in TB patients exhibited marked reductions in IgM positivity – reflecting a global, systemic shift in the B cell compartment – while B:M_14-_ doublets specifically displayed decreased IgD positivity compared to healthy controls (**Figure S5D**). For CD14^+^ monocyte complexes, both singlets and T:M doublets showed increased CD16 positivity in TB patients, indicating an enrichment for intermediate monocytes (**Figure S5F**). Thus, surface immunophenotyping of circulating doublets yielded minimal disease-specific changes beyond those already evident in the singlet pool.

In contrast, high-dimensional imaging feature extraction uncovered significant, disease-specific morphometric alterations. For CD3 imaging parameters, diffusivity and short-axis moment were significantly elevated across all T cell doublet types and singlets in TB patients, whereas maximum intensity was broadly reduced (**Figure 6A**). Doublet-exclusive morphometric shifts included increased long-axis moment in T:M and T:M_14-_ complexes, an increased radial moment in T:M and T:T doublets, and decreased total intensity in T:T doublets (**Figure 6A**). For CD14 imaging features, diffusivity and short-axis moments were consistently increased across all CD14^+^ monocyte doublet types and singlets in TB patients (**Figure 6B**). Radial moment was uniquely elevated in M:B, M:NK, and M:M doublets, whereas long axis moment was increased specifically in M:B and M:M pairs (**Figure 6B**). CD19 imaging parameters exhibited the fewest disease-associated alterations, with TB marked by increased short-axis moment in B:M and B:M_14-_ doublets and elevated radial moment in B:M and B:B doublets (**Figure 6C**). Together, these results demonstrate that imaging-derived parameters reveal disease-specific alterations in cell doublet morphology that are not detected by conventional fluorescence profiling.

**Figure 6:**
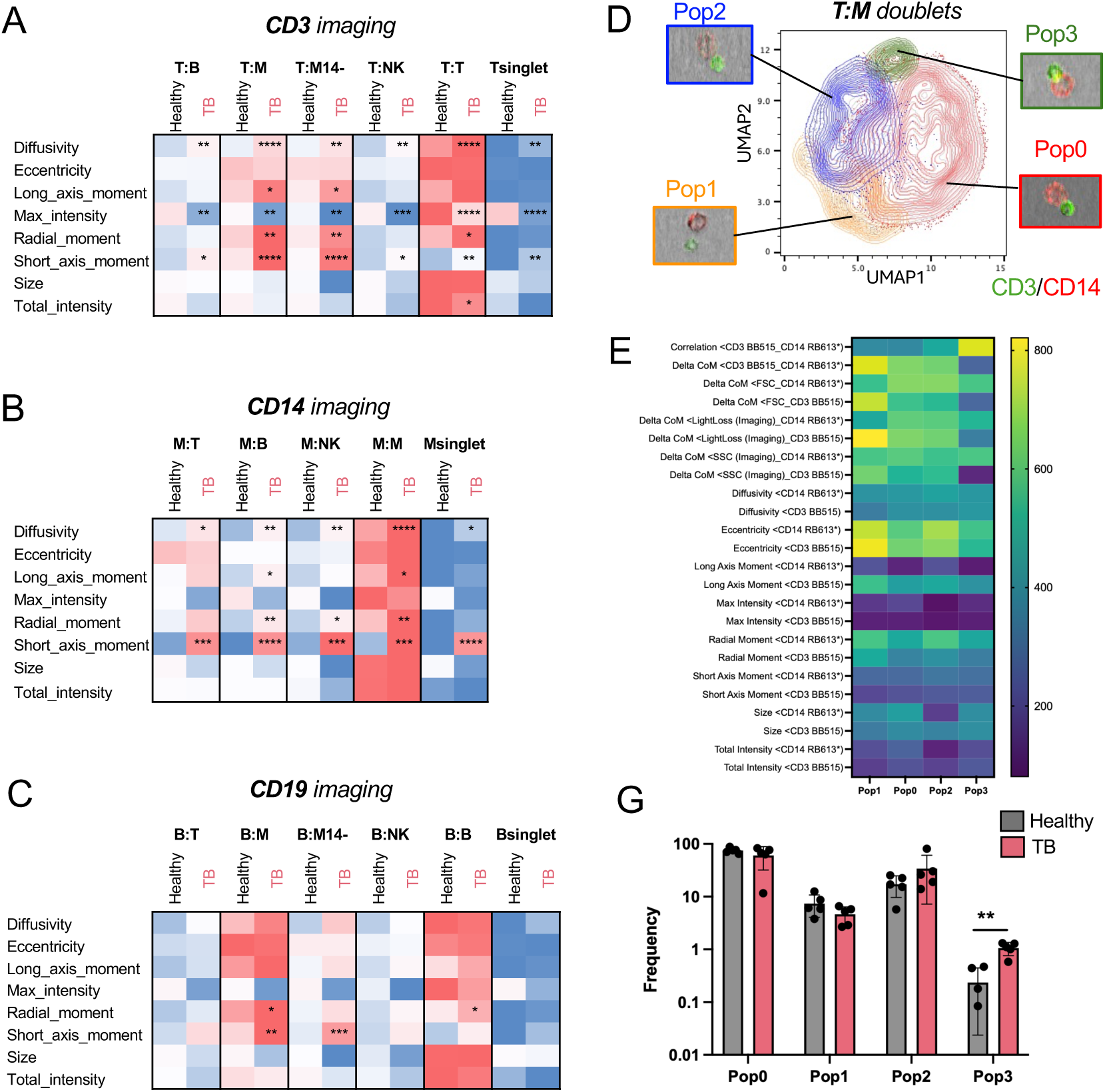
Quantitative imaging parameters uncover disease-specific doublet morphologies in active tuberculosis. Heatmaps depicting normalized median values of (A) CD3 imaging parameters for T cell-containing, (B) CD14 imaging parameters for CD14^+^ monocyte-containing, and (C) CD19 imaging parameters for B cell-containing doublets and singlets, in healthy controls versus TB disease patients (blue, low; red, high). (D) Unsupervised UMAP clustering and representative images of CD3 and CD14 imaging parameters in T cell-monocyte doublets across combined cohorts. (E) Heatmap of normalized median imaging features across UMAP clusters. (F) Relative cluster abundance (expressed as a percentage of total T cell-monocyte doublets) in healthy donors (n=5) versus TB disease patients (n=5). Individual datapoints, median and interquartile range are plotted. Cohorts were compared using an unpaired Mann-Whitney test (** *p*<0.01). Image data were processed using FlowJo^TM^ (v10.10) and the BD CellView^TM^ Lens plugin (v2.1).

### Unsupervised clustering of imaging parameters uncovers close-contact T cell-monocyte synapses in TB disease

To further analyze imaging parameter heterogeneity within doublet populations and disease cohorts, we performed unsupervised UMAP clustering on two heterotypic configurations possessing dual-lineage imaging signals: T:M and T:B doublets. This approach enabled the evaluation of cross-channel imaging parameters, including fluorescence pixel correlation and delta center of mass (defined as the physical distance between the geometric centers of two opposing fluorescent signals). Because of the signal spillover of CD14 expression into the CD19-dedicated ImgB3 channel (**Figures 1F, 1G**), M:B doublets were excluded from this analysis.

Within the T:M doublet space, UMAP analysis resolved four distinct clusters (Pop 1-4, **Figure 6D**). Pop 3, the smallest cluster, exhibited high CD3/CD14 pixel correlation and minimal delta center of mass, denoting intimate cell-cell contact (**Figures 6E**, **S6A**). Conversely, Pop 1 was defined by high eccentricity and a large delta center of mass distance between CD3 and CD14, reflecting greater spatial separation between the two cells (**Figures 6E**, **S6A**). The remaining two populations comprised physically interacting pairs with a small contact area and notable differences in CD14 expression: Pop 0 exhibited bright CD14 expression, whereas Pop 2 displayed dimmer, more diffuse CD14 expression.

Strikingly, comparative clinical cohort analysis revealed that Pop 3 was significantly enriched in patients with TB compared to healthy controls (**Figure 6F**), indicating an increased frequency of tightly adhered T:M events during active disease. Equivalent UMAP analyses of the T:B doublet population revealed comparable architecture – including a high-correlation, tightly synapsed cluster (Pop 2 **Figures S6B**, **S6C**) – yet its relative abundance remained unchanged between healthy donors and TB patients (**Figures S6D**). In summary, although overall circulating doublet composition remains constant in TB, high-content imaging parameters successfully resolve disease-specific sub-phenotypes – specifically, an expansion of tightly synapsed T cell-CD14^+^ monocyte complexes.

## Discussion

In this study, we established an image-guided spectral sorting pipeline on the BD FACSDiscover™ S8 platform to systematically construct the first comprehensive atlas of circulating immune cell doublets in human peripheral blood. A key technical challenge in designing our 22-color panel was navigating optical spillover between fluorescent imaging channels in the absence of native image unmixing algorithms in earlier software versions (BD FACSChorus™ v5.4). While this limitation required us to exclude cross-channel imaging parameters between CD14 and CD19 (such as correlation, and delta center of mass), single-channel morphometrics and fluorescence remained unaffected. Crucially, recent updates to the FACSChorus™ software suite now incorporate full spectral unmixing across all imaging channels, thus effectively eliminating these optical constraints. However, it is important to note that the system permits unmixing exclusively *within* the dedicated imaging channels. Panel design must therefore continue to prevent spillover from non-imaging fluorochromes into imaging detectors.

By integrating marker-independent morphometric parameters (Side Scatter and LightLoss imaging) with targeted cell-surface markers, our workflow successfully purged coincidental transits and a substantial background of platelet-associated events. In conventional flow cytometry and droplet-based single-cell sequencing workflows, these CD42b⁺ platelet-leukocyte aggregates frequently mask true biological doublets. By systematically filtering out these contaminants, we isolated 12 highly pure, validated homotypic and heterotypic immune doublet configurations. A major insight from this panel design was that non-imaging surface markers must be strictly lineage-exclusive (e.g., CD11c versus HLA-DR for myeloid cells) to unambiguously determine which interacting partner expresses a given protein. While this requirement prevents the sub-phenotyping of homotypic doublets using non-imaging detectors – where cell-of-origin attribution remains impossible – this constraint can be overcome by placing the marker of interest directly into an imaging channel. Moving forward, this atlas framework is readily adaptable to interrogate other specialized cell populations; for instance, incorporating mutually exclusive markers such as CD1c or FcεRIα will enable fine-grained resolution of dendritic cell-containing complexes.

Evaluating these 12 doublet populations at steady state revealed that circulating cell-cell complexes do not form through random associations driven purely by cellular abundance. By normalizing doublet frequencies against precursor singlet abundances to calculate constants of association (Ka), we decoupled cellular prevalence from affinity. This classification demonstrated that while T cell-containing pairs dominate total abundance due to high singlet T cell numbers in the circulation, heterotypic doublets containing CD14⁻ myeloid cells exhibit the highest intrinsic affinities toward B and NK cells. Future studies leveraging image-guided sorting will be instrumental in physically validating these interaction strengths – for instance, by sorting pure doublet populations and re-acquiring them under increasing fluidic shear pressures to directly test whether mathematical Ka values correlate with physical bond strength.

Furthermore, surface marker profiling revealed that physical cell-cell engagement is governed by specific immune sub-lineages, rather than representing random pairings of the singlet pool. Heterotypic doublets selected for unique phenotypic states, evidenced by the enrichment of CD20^-^ B cells within B:M and B:M_14-_ doublets, the increased prevalence of CD16⁺ intermediate monocytes in M:B doublets, and the underrepresentation of TCRγδ⁺ T cells in T:M_14-_ complexes. These phenotypic preferences may point to specialized functional interactions in circulation; for instance, while physical contact and antigen presentation between macrophages and B cells are well-documented in secondary lymphoid organs (16,17), direct physical conjugation between myeloid cells and B cells in human peripheral blood has rarely been quantified or functionally characterized. Together, these findings indicate that distinct immune cell subsets are intrinsically predisposed to form circulating physical interactions, and specific pairing combinations likely execute distinct immunological functions.

Importantly, our comparative evaluation of healthy controls and active TB patients highlighted a major paradigm shift: quantitative imaging features derived from CD3, CD14 and CD19 lineage markers captured subtle disease-specific alterations that are entirely missed by conventional fluorescence expression or population frequency counting. While doublet abundances, Ka affinities, and surface marker profiles remained largely indistinguishable between healthy and TB cohorts, quantitative imaging features (such as CD3 and CD14 radial and long-axis moments) revealed pronounced divergence in specific subsets of doublets during TB disease that were not observed in singlets. This demonstrates that cell lineage marker morphometrics are far more sensitive readouts of systemic immune activation and physical cell-cell interactions than fluorescence expression alone.

Finally, unsupervised UMAP clustering of interactive spatial parameters unlocked hidden, disease-specific sub-phenotypes within T cell-monocyte doublets. We identified a distinct structural cluster (Pop 3) defined by high CD3/CD14 spatial pixel correlation and minimal delta center of mass distance, representing tightly synapsed, intimate physical complexes. The significant enrichment of this tightly synapsed cluster in patients with TB disease likely captures in vivo immunological synapse formation during ongoing systemic immune system activation. Together, this work establishes a robust, image-guided paradigm for dissecting genuine cell-cell communication, offering a powerful template for mapping physical immune networks in human health and disease.

## Material and methods

### Ethics statement

Human study participants were enrolled at the field site of the South African Tuberculosis Vaccine Initiative (SATVI; University of Cape Town, Western Cape Province, South Africa). Ethical approval to carry out this work was granted and maintained by the Institutional Review Board of the La Jolla Institute for Immunology and the Human Research Ethics Committee of the University of Cape Town for the Protection of Human Subjects. All clinical investigations were conducted according to the principles expressed in the Declaration of Helsinki, and all participants (or guardians for participants <18 years old) provided written, informed consent before participation in the study.

### Study cohorts and samples

TB disease was defined by: 1) the presence of clinical symptoms and/or radiological/histological evidence of pulmonary TB, and 2) microbiological confirmation via *Mycobacterium tuberculosis* (Mtb)-specific molecular or culture testing. Healthy controls had no past medical history of TB, no known exposure to Mtb and no evidence of Mtb sensitization as confirmed by a negative Interferon-Gamma Release Assay (IGRA). PBMC were isolated from whole blood samples by density gradient centrifugation using Ficoll-Paque^®^ PLUS (Cytiva) according to the manufacturer’s instructions. Isolated PBMC were resuspended at up to 10 million cells per milliliter in fetal bovine serum (FBS; Gemini Bio) containing 10% dimethyl sulfoxide (DMSO; Sigma-Aldrich) and cryopreserved in liquid nitrogen.

### PBMC thawing

Cryopreserved PBMC were rapidly thawed in a 37°C water bath for 2 minutes and immediately transferred into 9 ml of cold complete medium consisting of RPMI 1640 with L-Glutamine and 25 mM Hepes (Omega Scientific), supplemented with 5% human AB serum (GemCell), 1% penicillin/streptomycin (Gibco), 1% GlutaMAX (Gibco), and 20 U/mL Benzonase^®^ Nuclease (Millipore). Cells were centrifuged, resuspended in complete medium, and evaluated for cell concentration and viability using trypan blue exclusion and a hematocytometer. PBMC were maintained at 4°C prior to downstream antibody staining and acquisition.

### Ex vivo antibody staining

For surface immunophenotyping, up to 2 × 10^6^ thawed PBMC were resuspended in 70 μl of phosphate-buffered saline (PBS) containing 5 μl of Human BD Fc Block^TM^ (BD Biosciences) and incubated for 10 min at room temperature. All fluorochrome-conjugated primary antibodies were centrifuged at 10,000g to pellet aggregates, for 5 minutes at 4°C prior to use to pellet aggregates. Predetermined optimal volumes of fluorochrome-conjugated anti-human antibodies and viability dye (**Table 1**), along with 10 μl of Brilliant Stain Buffer (BD Biosciences) were added to each sample and incubated for 30 min at room temperature in the dark. Single-stained cellular controls and SpectraComp (SlingshotBio^TM)^ cell mimic controls were prepared concurrently using identical staining protocols and volumes. Following incubation, cells were washed twice in flow cytometry buffer (PBS containing 2 mM EDTA [pH 8.0] and 0.5% bovine serum albumin [BSA]), resuspended in 100 μl of flow cytometry buffer and stored at 4°C protected from light for up to four hours prior to acquisition.

### Flow cytometry data acquisition

Samples were acquired on a BD FACSDiscover^TM^ S8 Cell Sorter with BD SpectralFX^TM^ and BD CellView^TM^ Image Technology with the standard 5-laser configuration (349 nm, 405 nm, 488 nm, 561 nm, 637 nm), 6 imaging detectors (3 scatter, 3 fluorescence PMTs) and a spectral array of 78 APD detectors, operating on BD FACSChorus^TM^ software version 5.4. The 85μm nozzle with standard settings (35 psi, 57kHz) was used, and instrument and imaging calibration QC were performed using BD FACSDiscover^TM^ Setup Beads and BD CellView^TM^ Calibration Beads, respectively. Gain settings for LightLoss (Violet) and SSC were adjusted to optimize resolution of immune cells within PBMC for the primary gate. QC-determined gain settings were used for all other detectors. An unstained sample, single-stained cell and cell mimic controls were acquired and used to optimize the unmixing matrix applied to all samples. For imaging optimization, the Region of Analysis (ROA) was adjusted based on SSC, then Pixel Threshold for each imaging detector was set appropriately, followed by adjustments to channel settings to minimize overall background. All samples were acquired at <8,000 events/sec and FCS files with and without images were recorded.

### Flow cytometry data analysis

High-dimensional fluorescence and imaging data were analyzed using FlowJo^TM^ software (version 10.10). Imaging data was visualized either natively within BD FACSChorus™ (ImageWall) or via the BD CellView™ Lens plugin (v2.1), as indicated. Staining index (SI) was calculated in FlowJo^TM^ (version 10.10) as (MFI_positive − MFI_negative) / (2 × SD_negative), as previous described (18). Statistical analyses and graphing were performed using GraphPad Prism (version 10.4). Pairwise comparisons between doublet configurations and singlet counterparts, as well as between healthy and TB disease cohorts, were evaluated using two-tailed, non-parametric Mann–Whitney U tests. P-values less than 0.05 were considered statistically significant. Principal Component Analysis (PCA) was performed in R (version 4.5) using the prcomp function on centered and scaled morphometric datasets. Unsupervised clustering and UMAP dimensional reduction were performed in FlowJo™ (version 10.10) using the FlowSOM (version 3.0) and UMAP (version 3.1) plugins.

