## Supplemental Figures for "A high-resolution blood immune cell doublet atlas via imaging spectral cytometry"

<u>Content</u>

*Figures S1 to S6*

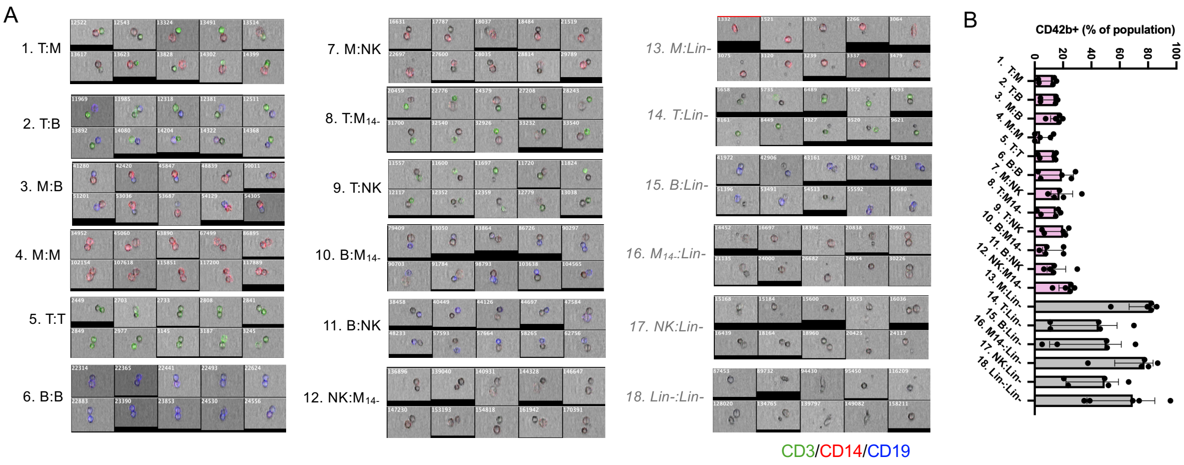


**Figure S1: Lineage-negative doublet gates harbor substantial platelet contamination.** (A) Representative composite image galleries comparing the 12 validated immune doublet populations to the 6 ambiguous lineage-negative doublet populations identified in Figure 2C (CD3, green; CD14: red; CD19: blue). (B) Frequency of CD42b$^{+}$ platelet-associated events within the 12 main doublet populations (pink) and the 6 lineage-negative doublet populations (grey). Individual datapoints, median and interquartile range are plotted. Data derived from healthy PBMC (n=5).

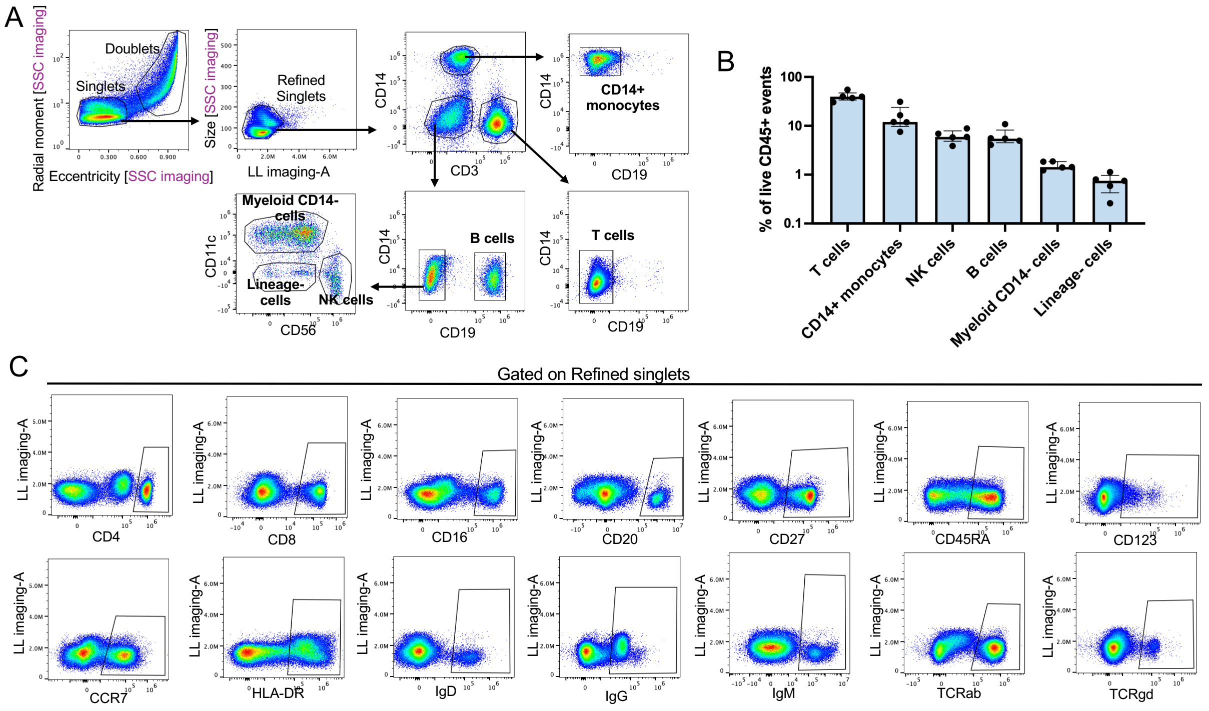


**Figure S2: Gating strategy, frequency and surface phenotype of circulating singlets.** (A) Gating strategy resolving singlet T cells (CD3$^{+}$CD14$^{-}$CD19$^{-}$), CD14+ monocytes (CD3$^{-}$CD14$^{+}$CD19$^{-}$), B cells (CD3$^{-}$CD14$^{-}$CD19$^{+}$), CD14$^{-}$ myeloid cells (CD3$^{-}$CD14$^{-}$CD19$^{-}$CD11c$^{+}$CD56$^{-}$), NK cells (CD3$^{-}$CD14$^{-}$CD19$^{-}$CD11c$^{-}$CD56$^{+}$), and lineage-negative cells (CD3$^{-}$CD14$^{-}$CD19$^{-}$CD11c$^{-}$CD56$^{-}$) from human PBMC. (B) Frequencies of singlet populations expressed as a percentage of live CD45$^{+}$ events. Individual datapoints, median and interquartile range are plotted. (C) Representative staining profiles for the 14 non-gating phenotypic markers in refined singlets. Individual datapoints, median and interquartile range are plotted. Data derived from healthy PBMC (n=5).

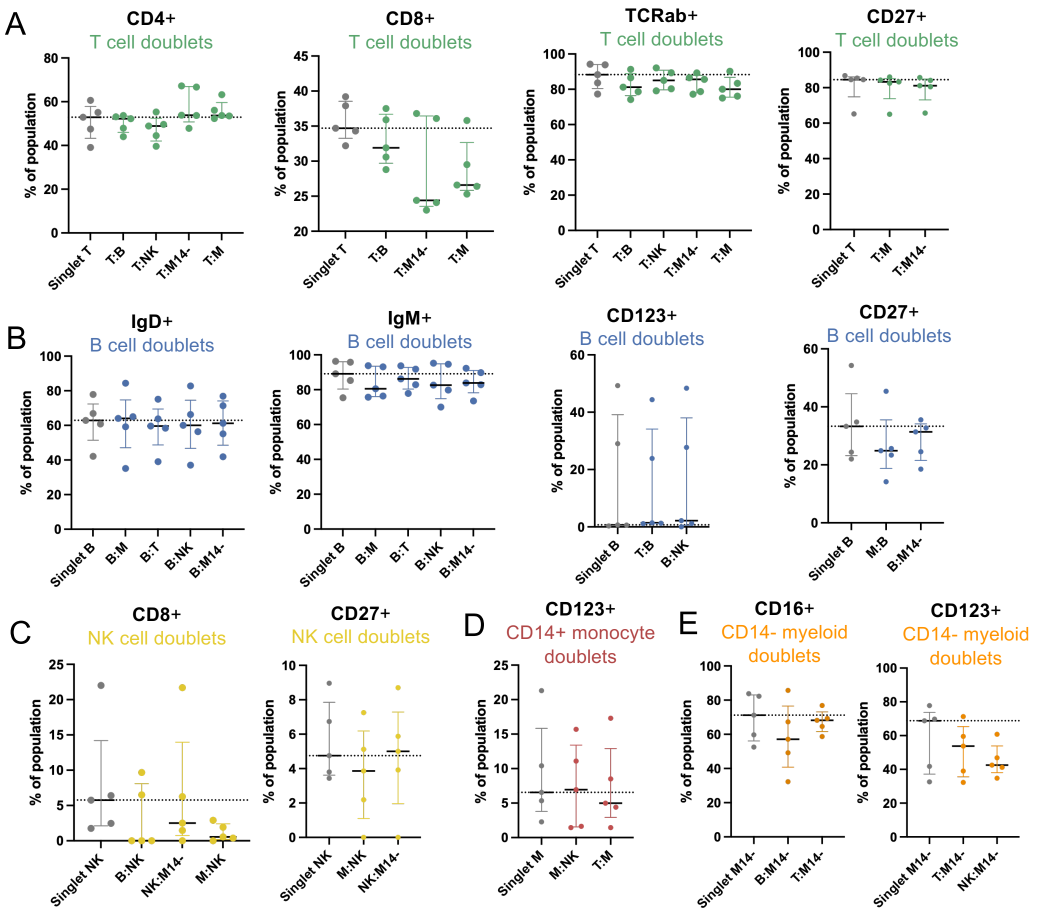


**Figure S3: Surface phenotypes that do not significantly differ between doublet populations and singlets.** Frequencies of marker positive cells across (A) T cell-, (B) B cell-, (C) NK cell-, (D) CD14+ monocyte-, and (E) CD14- myeloid cell-containing heterotypic doublet configurations compared to singlet counterparts. Doublet populations were identified as shown in Figure 2C. Individual datapoints, median and interquartile range are plotted; dashed line represent median frequency of singlets. Frequencies of marker-positive cells within heterotypic doublet populations were compared to singlets using a non-parametric unpaired Mann-Whitney test. Data derived from healthy PBMC (n=5).

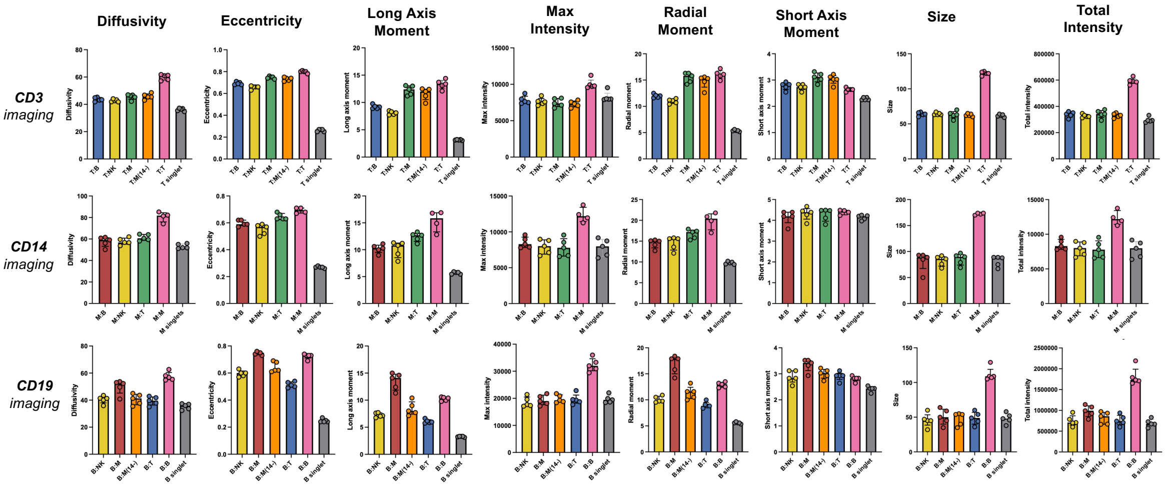


**Figure S4: Imaging features from CD3, CD14, and CD19 channels differ between doublet pairings.** Quantitative distribution of CD3 (top), CD14 (middle), and CD19 (bottom) imaging parameters across indicated T cell-, $CD14^+$ monocyte-, and B cell-containing doublet populations compared to singlet counterparts. Doublet populations were identified as shown in Figure 2C and singlets as shown in Figure S2A. Individual datapoints, median and interquartile range are plotted. Data derived from healthy PBMC (n=5).

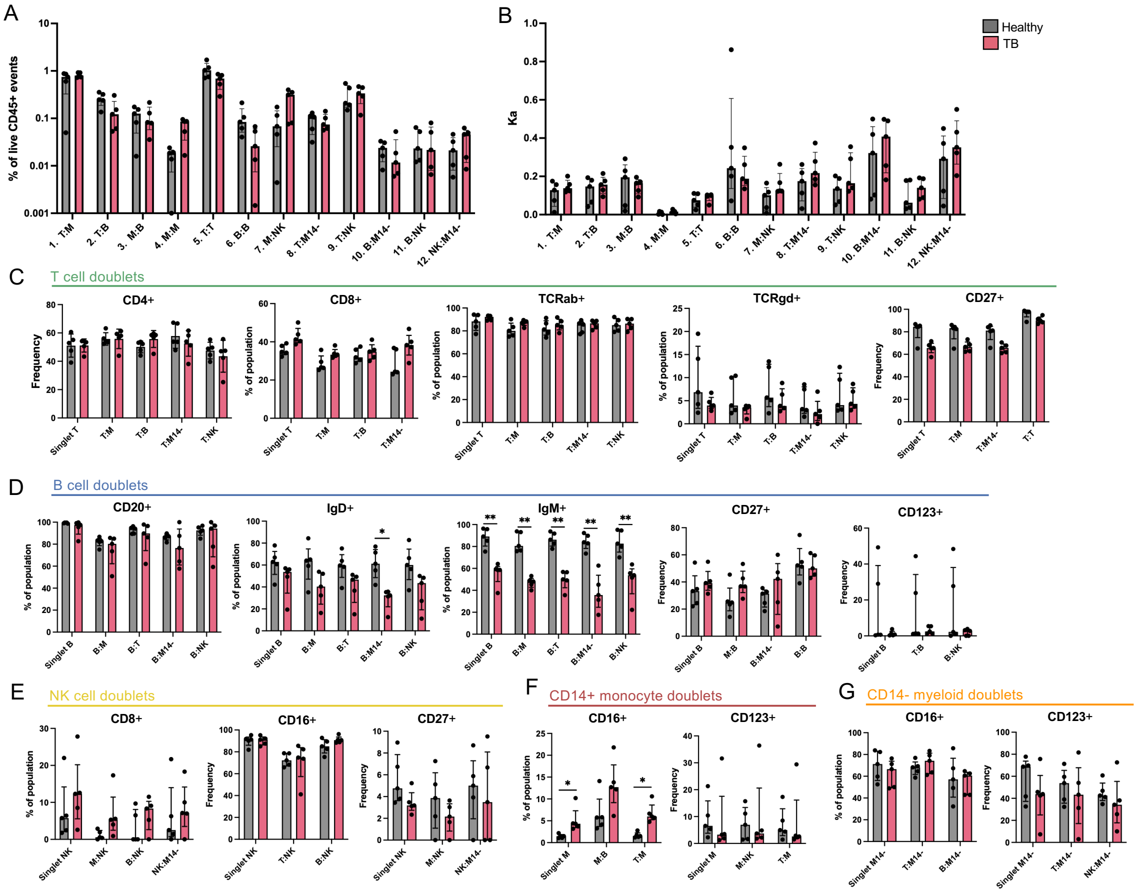


**Figure S5: Doublet frequencies or surface phenotypes do not differ between healthy and TB cohorts.** (A) Frequencies (expressed as a percentage of live $CD45^{+}$ events) and (B) constant of association Ka of the 12 validated doublet populations in healthy (grey, n=5) and TB disease (pink, n=5) cohorts. Ka was calculated by dividing doublet frequency by the product of constituent singlet frequencies, as previously reported (2). Frequencies of marker positive cells across (C) T cell-, (D) B cell-, (E) NK cell-, (F) CD14+ monocyte-, and (G) CD14- myeloid cell-containing indicated heterotypic doublets and singlet counterparts between healthy and TB cohorts. Doublet populations were identified as shown in Figure 2C and singlets as shown in Figure S2A. Individual datapoints, median and interquartile range are plotted. Cohorts were compared using a non-parametric unpaired Mann-Whitney test (** $p<0.01$).

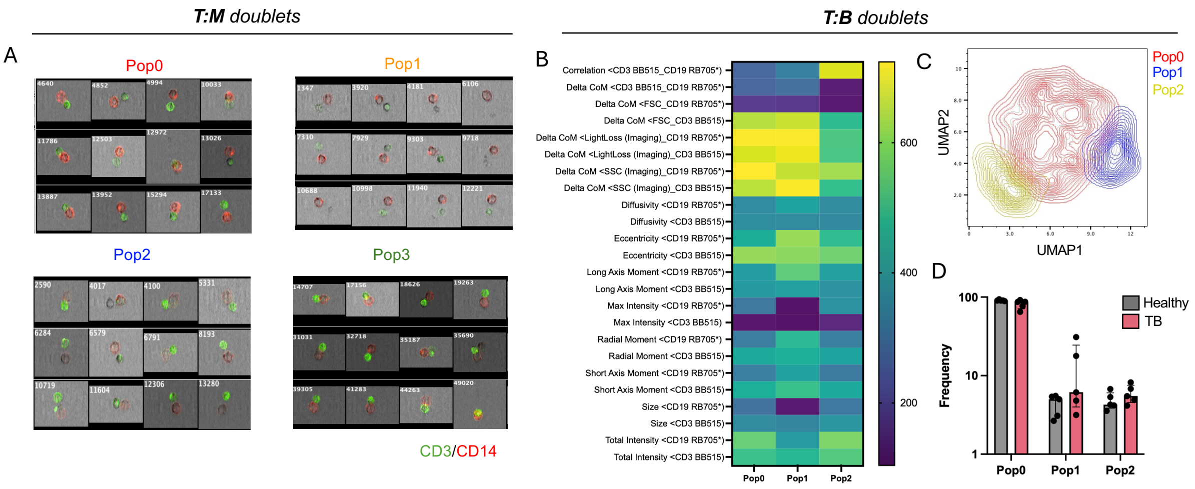


**Figure S6: Unsupervised clustering of imaging parameters in T cell-monocyte and T cell-B cell complexes.** (A) Representative image galleries across the four UMAP clusters identified from unsupervised clustering of CD3 (green) and CD14 (red) imaging parameters in T cell-monocyte complexes (Figure 6D). (B) Unsupervised UMAP clustering and representative images of CD3 and CD19 imaging parameters in T cell-B cell doublets across combined cohorts. (C) Heatmap of normalized median imaging features across UMAP clusters. (D) Relative cluster abundance (expressed as a percentage of total T cell-B cell doublets) in healthy donors (n=5) versus TB disease patients (n=5). Individual datapoints, median and interquartile range are plotted. Cohorts were compared using an unpaired Mann-Whitney test. Image data were processed using FlowJo$^{TM}$ (v10.10) and the BD CellView$^{TM}$ Lens plugin (v2.1).
